# Experimental sexual selection affects the evolution of physiological and life history traits

**DOI:** 10.1101/2021.03.23.436586

**Authors:** Martin D. Garlovsky, Luke Holman, Andrew L. Brooks, Zorana K. Novicic, Rhonda R. Snook

**Affiliations:** Department of Animal and Plant Sciences, University of Sheffield, UK; Applied Zoology, Faculty Biology, Technische Universität Dresden, Germany; School of Applied Sciences, Edinburgh Napier University, UK; Animal Ecology, Department of Ecology and Genetics, Evolutionary Biology Center, Uppsala University, Uppsala, Sweden; Department of Zoology, Stockholm University, Sweden

**Author notes:** These authors contributed equally to this work (Martin D. Garlovsky, Luke Holman).

**Keywords:** Sexual selection, polyandry, metabolism, physiology, life history evolution, trade-offs, experimental evolution

## Abstract

Sexual selection and sexual conflict are expected to affect all aspects of the phenotype, not only traits that are directly involved in reproduction. Here, we show coordinated evolution of multiple physiological and life history traits in response to long-term experimental manipulation of the mating system in populations of *Drosophila pseudoobscura*. Development time was extended under polyandry relative to monogamy in both sexes, potentially due to higher investment in traits linked to sexual selection and sexual conflict. Individuals (especially males) evolving under polyandry had higher metabolic rates and locomotor activity than those evolving under monogamy. Polyandry individuals also invested more in metabolites associated with increased endurance capacity and efficient energy metabolism and regulation, namely lipid and glycogen. Finally, polyandry males were less desiccation- and starvation-resistant than monogamy males, suggesting trade-offs between resistance and sexually selected traits. Our results provide experimental evidence that mating systems can impose selection that influences the evolution of non-sexual phenotypes such as development, activity, metabolism, and nutrient homeostasis.

## INTRODUCTION

Natural and sexual selection often act differently in males and females, which have different routes to evolutionary fitness as a result of anisogamy (Darwin, 1871; Kokko & Jennions, 2008). However, independent evolution of the sexes is partly constrained due to their shared genome (Lande, 1980). Recent work has increasingly highlighted that sexual selection affects all aspects of the phenotype, and not only classic sexually-selected traits such as male ornaments (reviewed in Cally *et al*., 2019). This is because sexually-selected traits often depend on overall condition (Rowe & Houle, 1996), which in turn depends on underlying aspects of organisms such as their development, physiology, life history and metabolism (Lailvaux & Irschick, 2006; Orr & Garland, 2017).

Sexual selection often favours energetically costly bouts of sustained locomotor activity, e.g. during mate searching, courtship, or the ‘harassment and resistance’ behaviours that typify interlocus sexual conflict (Watson *et al*., 1998; Kotiaho, 2001; Hunt *et al*., 2004; Gyulavári *et al*., 2014; Debelle *et al*., 2017). Moreover, postcopulatory sexual selection can select for males that produce metabolically expensive ejaculates (Linklater *et al*., 2007; Immonen *et al*., 2016). We therefore expect sexual selection and sexual conflict to favour physiological adaptations that augment the efficiency of metabolism and respiration (Montooth *et al*., 2003; Gyulavári *et al*., 2014). Sexual selection is also hypothesised to affect the evolution of development and life history, potentially in a sex-specific way (Badyaev, 2002; Stångberg *et al*., 2020). For instance, in populations experiencing heightened sexual selection, male *Drosophila melanogaster* evolved increased activity and courtship frequency and shorter lifespan (Nandy *et al*., 2013), and in a separate experiment, males experiencing heightened sexual selection evolved faster development time, whereas females developed faster under monogamy (Hollis *et al*., 2017). Likewise, selection for early life reproduction in *Acanthoscelides obtectus* beetles favoured higher metabolic rate in males (Arnqvist *et al*., 2017), and in *Callosobruchus maculatus* beetles, females (but not males) evolved under polygamy had higher mortality and aging rates than females evolved under monogamy (Maklakov *et al*., 2007). Thus, selection may cause trade-offs between different aspects of fitness, for instance, favouring a ‘live fast, die young’ strategy in species where competition for mates is intense (Nandy *et al*., 2013; Hollis *et al*., 2017; Hämäläinen *et al*., 2018) or selecting for the reallocation of limiting resources away from growth and somatic maintenance (Pitnick *et al*., 1995; Hunt *et al*., 2004; Emlen *et al*., 2012; Berson *et al*., 2019). Indeed, the conspicuous fitness trade-offs associated with sexually-selected traits was a key motivation for the theory of sexual selection (Darwin, 1871).

In addition to these links between sexually selected traits and other phenotypes, the picture is further complicated by genetic correlations between the sexes. Male and female traits have a shared genetic basis, such that selection on males results in a (frequently maladaptive) evolutionary response in females, and *vice versa* e.g. (Long & Rice, 2007; Harano *et al*., 2010; Lewis *et al*., 2011; Berger *et al*., 2014; Jensen *et al*., 2015; Arnqvist *et al*., 2017; Holman & Jacomb, 2017; Hämäläinen *et al*., 2018; Immonen *et al*., 2018; Videlier *et al*., 2019). Therefore, sexual selection on males produces an evolutionary response in female, as well as male, life history traits and vice versa. For instance, male-limited selection for short life span in *C. maculatus* caused a correlated response in females (Berger *et al*., 2014), and in *D. melanogaster*, male-limited evolution selected for increased activity in both sexes, to the detriment of female fitness (Long & Rice, 2007). While these studies have measured the response of individual or several key physiological or life history traits to the strength of sexual selection, in this paper, we investigate the effect of variation in the strength of sexual selection on multiple inter-related life history and physiological traits to capture a broader range of consequences of sexual selection.

We use the fruit fly, *Drosophila pseudoobscura*, subjected to either experimentally enforced monogamy (M) that eliminated sexual selection and sexual conflict, or to elevated polyandry (E), where 6 males were housed with 1 female to facilitate inter- and intra-sexual selection, likely at elevated levels compared to natural populations (Anderson, 1974). For each sexual selection treatment, there were four replicate populations (hereafter ‘lines’). Previous studies have found divergence between the E and M lines in several traits important during episodes of sexual selection. For example, E males produce more abundant and complex chemical signals (Hunt *et al*., 2012), perform a faster and more vigorous courtship song (Snook *et al*., 2005; Debelle *et al*., 2014, 2017), court and mate more frequently (Crudgington *et al*., 2010), are more competitive in mating encounters (Debelle *et al*., 2016), and have larger glands for producing seminal fluid (Crudgington *et al*., 2009). Additionally, coevolutionary patterns have been found, such that E males are more harmful but E females more resilient to such harm (Crudgington *et al*., 2005, 2009), and females prefer courtship song of males from their own treatment (Debelle *et al*., 2014). These evolved phenotypic differences are accompanied by divergence in gene expression (Immonen *et al*., 2014; Veltsos *et al*., 2017) and concerted genetic differences between treatments (Wiberg *et al*., 2021).

Given these evolved differences in traits directly associated with reproduction, we tested whether the mating system similarly caused divergence in life history and physiological phenotypes that likely contribute to reproductive success, namely metabolic rate, macro-metabolite content, development time, and stress resistance. Assuming that these traits contribute to reproductive success, we predict that they may have diverged between the E and M lines, reflecting the different behaviours, life history decisions, and physiological states that maximise fitness under the different selective regimes. Due to the genetic non-independence of the sexes, the multiple differences in selection on both sexes selection in the E and M treatments, and the genetic correlations between the variables examined, the direction of evolutionary change is difficult to predict, making our study exploratory in nature. However, we predict higher activity levels and metabolic rates in the E treatment in both sexes, due to the elevated importance of courtship, harassment, and resistance behaviours. As such, elevated metabolism might require altered resource allocation, affecting macrometabolite composition, development time, and the ability to resist stressors. Shorter development may be favoured by sexual selection (Hollis *et al*., 2017), however the effects of sexual selection on macrometabolite profile is unknown *a priori*.

## METHODS

### Establishment and maintenance of experimental evolution lines

Details of the establishment and maintenance of the experimental evolution lines has been previously described (Crudgington *et al*., 2005). Briefly, the ancestral population was established from 50 wild-caught, inseminated female *Drosophila pseudoobscura* from Tucson, Arizona in 2001; the descendants of these females were used to establish four replicate populations (termed lines) for each sexual selection treatment. In the enforced monogamy or ‘M’ treatment, the adult population was housed in groups of two (one male, one female); this protocol reduces the opportunity for sexual selection and sexual conflict. In the elevated polyandry or ‘E’ treatment, each group of adults comprised six males and one female. Each M line contained a greater number of groups (n = 80) than each E line (n = 40), to offset the smaller group size and thereby minimise differences between treatments in the autosomal effective population size (Snook *et al*., 2009). In each generation, unmated males and females were housed in ‘interaction vials’ (IVs) for 5 days, before being transferred to ‘oviposition vials’ (OVs) for a further 5 days; using both IVs and OVs reduces larval competition and provides more opportunity for episodes of pre- and post-copulatory sexual selection. To facilitate selection favouring the most productive groups (and the most competitive males, for the E treatment), offspring from all groups within each line were gathered and mixed in a single container, and a random sample of all the offspring was used to set up the next generation. Flies were kept at 22ºC on a 12:12 light:dark cycle on standard cornmeal-agar-molasses media with added live yeast.

### Experimental individuals

Prior to the experiments described below, flies from each line were taken out of selection and placed in a ‘common garden’ to minimise non-genetic differences between lines. Newly-eclosed individuals were collected *en masse* from the OVs within each line. A random sample of these flies were allowed to mate and oviposit for two days. From these eggs, we set up controlled density vials (CDVs) by placing 100 first instar larvae into food vials. Unmated flies eclosing from CDVs were collected and stored in same-sex food vials until 3-5 days old. Excluding the development time experiment each measurement in our study was performed on a ‘triad’ of three age-matched, same sex flies within each line, due to practical constraints resulting from the small size of individual flies.

### Juvenile development time

We measured juvenile development time at generations 180, 179, 178 and 176 for lines 1-4, respectively. For each replicate of the E and M treatments, we seeded 6 CDVs on three consecutive days (i.e., 600 larvae per replicate population per seeding day = 14,400 larvae).

Vials were checked daily for new eclosees, and flies were CO_2_ anaesthetised and killed in ethanol. We continued collecting until no individuals eclosed for two consecutive days. We subsequently counted the number of adult males and females emerging each day from each vial.

We used the length of wing vein four as a proxy for body size (Crudgington *et al*., 2005) by measuring a random subsample of individuals (n = 15 per sex per replicate per seeding day). We removed the left wing from flies preserved in ethanol and mounted wings on a microscope slide in a drop of phosphate-buffered saline and dried at room temperature overnight. We imaged wings using a Motic camera and Motic Images Plus 2.0 software (Motic Asia, Hong Kong). We measured wing vein four length using ImageJ software (Schneider *et al*., 2012). Image files were anonymised prior to measurement.

#### Desiccation and starvation resistance

We measured desiccation and starvation resistance at generations 199, 198, 197 and 195 for lines 1-4, respectively. Triads (n = 7-10) were housed in 8-dram plastic vials stoppered with cotton balls and covered with Parafilm^®^. For the desiccation resistance assay, vials contained no food and between the cotton and Parafilm^®^ we placed a packet of silica gel beads. For the starvation resistance assay, vials contained an agar solution that provided moisture but no food. Vials were checked every 2 hours and any deaths recorded until all flies perished. Flies were scored as dead if they were not able to right themselves or no movement was observed.

#### Respirometry

We measured metabolic rates at generations 196, 195, 194 and 192 for lines 1-4, respectively. Triads (n = 3) were weighed to the nearest 0.1 mg (Sartorius Genius ME 235P-OCE) before transfer to a respirometry chamber (a glass cylinder; 17mm × 70 mm). Metabolic rate was measured using a Sable Systems (Las Vegas, NV, USA) respirometry system (Lighton, 2008). This system pumps air at a precise flow rate through a sealed chamber containing the organisms undergoing measurement. Downstream gas analysers measure two response variables: the amount of CO_2_ produced and O_2_ consumed. Briefly, the respirometry system was set up in stop-flow mode (Lighton, 2008), in which each chamber was sealed for 60 min and then flushed for 2.5 min. Each cycle (through all 24 chambers) lasted for 62.5 minutes, and each triad of flies was recorded over consecutive cycles, giving four readings of CO_2_ and O_2_ flux in each individual chamber. The first recording was discarded as a wash-out, while the other three-were used for analyses. Each respirometry chamber was placed in an activity detector (AD-2, Sable Systems) connected to a data acquisition interface (Quick-DAQ, National Instruments, Coleman Technologies, Newton Square, US), which uses reflective infrared-light technology to provide a precise and continuous measure of locomotor activity of the subjects; one activity measurement per triad per cycle was recorded. One of the 24 chambers was left empty and used as a baseline to control for any drift of the gas analysers during each session (washed out twice in each cycle). Thus, each observation consisted of three consecutive readings of the amount of CO_2_ produced and O_2_ consumed during 62.5 minutes by a triad of flies, under dark conditions, with a known weight and total amount of activity performed. We also calculated the respiratory quotient (RQ; i.e., VCO_2_ / VO_2_), which describes the metabolic substrate used for respiration, where a value of 0.7 indicates pure fatty acid oxidation, 1.0 indicates pure carbohydrate oxidation, and intermediate values indicate mixed substrate or protein-based oxidation

### Metabolite extractions

We measured metabolite composition at generations 199, 198, 197 and 195 for lines 1-4, respectively. Triads (n = 3) were weighed to the nearest 1μg (METTLER TOLEDO^®^ UMX2 ultramicrobalance) and flash frozen in liquid nitrogen. Triads were then placed in a 0.35ml glass vial insert (SUPELCO Analytical^®^) of known weight, dried at 55ºC overnight, and re-weighed to obtain dry weight. We subsequently processed each sample to acquire measurements of lipids, soluble carbohydrates, soluble protein, glycogen, and chitin (see online supplementary material).

### Statistical analysis

All statistical analyses were performed in R version 4.0.3 (R Core Team, 2020). We fit models using Bayesian approaches in *brms* (Bürkner, 2017), and specified conservative, regularising priors on the fixed and random effects. Complete code and description of models are provided in the code repository (https://anonymous.4open.science/r/exp_evol_respiration-2647). In brief, we analysed development time and survival using survival analysis. For the respirometry and metabolite data we used structural equation models (SEM), motivated by the causal relationships that we hypothesise to exist between the variables, given the experimental design and our biological understanding of the system (Fig. S1). For the metabolite data, we began by expressing the abundance of each metabolite as a proportion of dry weight and scaled each response variable to have mean zero and unit standard deviation. For the SEMs, after fitting the model, we calculated the treatment effect size (Cohen’s *d*) for each response variable, both with and without the moderators (dry weight and/or activity). We also calculated the difference in treatment effect size between the sexes, to test for a treatment × sex interaction. All models included replicate line as a random intercept to reflect the experimental design of the selection experiment (i.e. where n = 8). We also performed analyses with selection treatment fitted as a random slope to allow lines to vary in their response to selection (Schielzeth & Forstmeier, 2009), however results were qualitatively identical, and thus we present only the results of models without random slopes in the Results. All models also included vial identity as a random intercept, except for wing vein length where we measured a random sample of individuals across multiple vials.

## RESULTS

### Juvenile development time

Juvenile development time differed significantly between the E and M treatments (Cox proportional hazards model; PP = 0.008, where PP is the posterior probability the effect is of the opposite direction, similar to a 1-tailed p-value) and between the sexes (PP < 0.001). E treatment flies had a reduced hazard (Hazard ratio = 0.44; 95% confidence intervals [CI] = 0.26 – 0.81), i.e., development took longer in the E treatment in both sexes. Males took longer to eclose than females (Hazard ratio = 0.84; 95% CI = 0.80 – 0.88) (Fig. 1A), as usual in this species. There was a significant treatment × sex interaction (PP = 0.007), such that the sex difference in development time was greater in the E treatment, and the treatment affected males more strongly than females.

**Figure 1:**
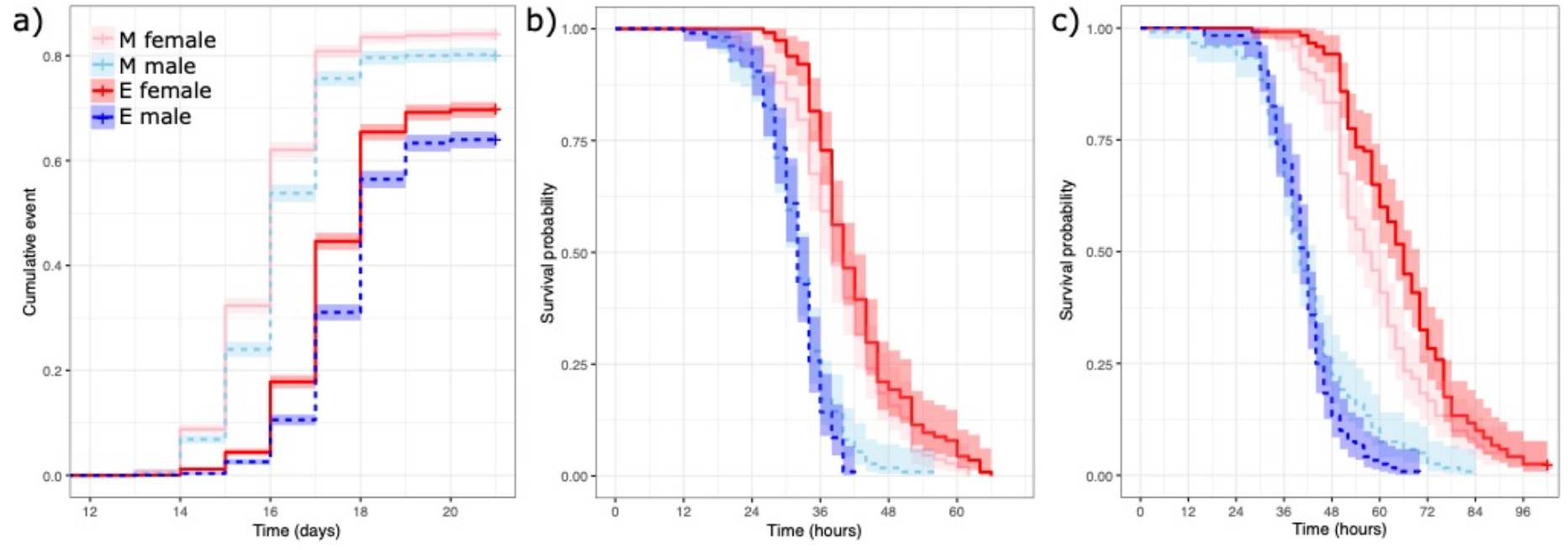
Kaplan-Meier plots for (a) juvenile development time, (b) desiccation resistance, and (c) starvation resistance. ‘+’ indicates right censored individuals. Monogamy (M) = pink and light blue, Elevated polyandry (E) = red and dark blue; Males = dashed lines, Females = solid lines. Shaded areas show confidence intervals.

As expected, females were larger than males (mean wing vein IV length (μm); M females: 2324 ± 4.80, n = 152; E females: 2335 ± 5.76, n = 118; M males: 2099 ± 4.93, n = 154; E males: 2114 ± 5.96, n = 127). We found no statistically significant effect of sexual selection treatment (PP = 0.312) or the treatment × sex interaction (PP = 0.382).

### Desiccation and starvation resistance

We found a significant treatment × sex interaction for both desiccation and starvation resistance (desiccation: PP = 0.024; starvation: PP = 0.018). E males survived 0.91 (95% CI = 0.83 – 0.99), and 0.88 (95% CI = 0.80 – 0.97), times as long as M males under desiccation and starvation resistance, respectively (Fig. 1B-C). However, the treatment effect was opposite in females: E females survived 1.06 (95% CI = 0.87 – 1.29), and 1.09 (95% CI = 0.88 – 1.35), times longer than M females under desiccation and starvation resistance, respectively. In short, M males survived longer than E males, while E females survived longer than M females.

### Respirometry

Beginning with the mediator variables, females were heavier than males (PP < 0.001; Table S1), and E females were non-significantly heavier than M females (Fig. 2A; Mean dry weight per triad (mg): M females: 0.40 ± 0.02; E females: 0.47 ± 0.04; M males: 0.29 ± 0.02; E males: 0.29 ± 0.02; n = 12 each). The treatment (PP = 0.075) and treatment × sex (PP = 0.105) terms were not significantly related to body mass. There was a significant effect of sexual selection treatment on activity level (PP < 0.001), with E flies more active than M flies (Fig. 2A). Activity level also showed a sex × cycle interaction (PP < 0.026), indicating that activity levels declined over cycles in males but not females. This sex difference in the rate of decline was stronger in M treatment than E treatment, though the 3-way interaction was not statistically significant (PP = 0.066).

**Figure 2:**
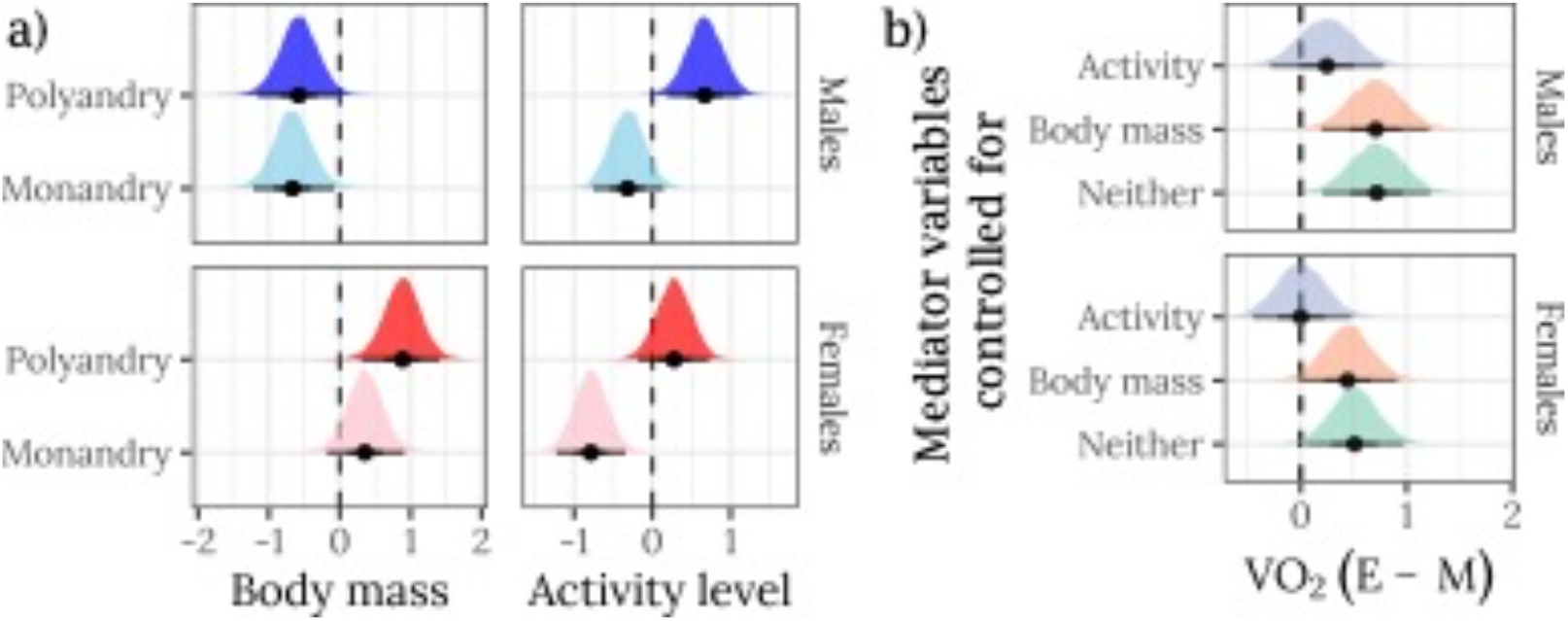
Effects of sex and selection treatment on metabolic rates and mediator variables. A) Posterior estimates of the means for mediator variables (body mass and activity). Note that females are larger than males, and that E females are somewhat larger than M females. B) Posterior estimates of effect sizes of selection treatment (E minus M) on metabolic rate (measured as volume of O_2_ consumed) controlling for activity, body mass, or neither mediator variable. The x-axes show the posterior estimate of standardised effect size (Cohen’s d), i.e., a value of 1 would mean that the E treatment has a mean that is larger by 1 standard deviation. The horizontal bars show the 66% and 95% quantiles and the median of the posterior distribution. All response variables have been mean-centred and divided by the overall standard deviation (such that the dashed line at zero marks the mean across sexes, treatments, and cycles). For brevity, only the first cycle is shown for activity and metabolic rate; see online supplementary material for all cycles.

There was a strong correlation between activity level and O_2_ consumption (PP < 0.001; Fig.S2), and O_2_ consumption declined across cycles (PP < 0.001). Body weight did not correlate with O_2_ consumption (PP = 0.119; Fig. S2). Calculating the posterior estimate of the difference in treatment means for each sex and cycle revealed that O_2_ consumption was higher in the E treatment (indicated by positive effect sizes in Fig. 2B), especially in males. This effect was considerably attenuated when we statistically adjusted for differences in activity level between treatments and sexes, indicating that the evolved difference in activity between the E and M treatments was largely responsible for the treatment effect on O_2_ consumption. Controlling for the evolved differences in body mass did not change the effect size, illustrating that the (modest) changes in body mass between the E and M treatments did not explain the evolved difference in O_2_ consumption.

The respiratory quotient (RQ) did not differ detectably between the E and M treatments, whether or not one controlled for the mediator variables (Fig. S3). The grand mean RQ = 0.90 (± 0.02) indicated flies used a mixture of carbohydrates, protein, and lipids as metabolic substrate.

### Metabolite composition

As in the respiration experiment, females were heavier than males and E flies were heavier than M flies (Dry weight (mg): M females: 0.56 ± 0.02; E females: 0.64 ± 0.17; M males: 0.33 ± 0.01; E males: 0.35 ± 0.01; n = 12 each; Table S2). There was a significant treatment × sex interaction (P = 0.038), such that the E treatment positively affected body mass more strongly in females. Furthermore, dry weight was significantly correlated with lipid and chitin content (Fig. S4; Table S2), and therefore could act as a mediator for some of the effect of sex and treatment on these metabolites.

We also found differences in metabolite composition between the E and M treatments and between sexes (Fig. 3). E treatment flies contained more lipids and glycogen than M flies, while M flies had relatively more carbohydrates and chitin; the amount of protein did not differ between treatments. These treatment differences reached statistical significance in males for carbohydrate, glycogen, and chitin, and in females for lipid. When we controlled for the evolved differences in dry weight between the E and M treatments, only the carbohydrate and glycogen differences between E and M males remained statistically significant; however, the actual change in effect size when controlling for dry weight was very small, indicating that dry weight was not an especially important mediator of the evolved changes in metabolite composition (compare Fig. 3 and Fig. S5).

**Figure 3.**
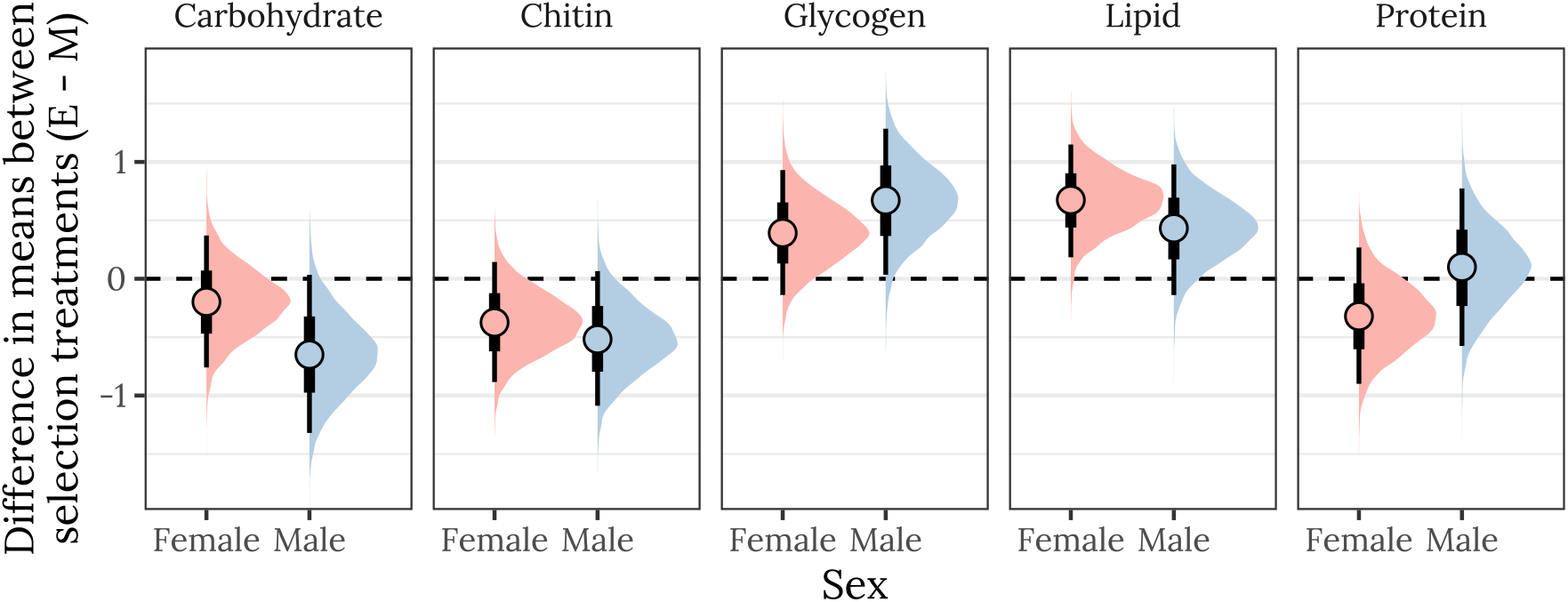
Posterior estimates of the treatment effect size for both sexes, for each of the five metabolites. A positive value indicates that the mean metabolite content is higher in the E treatment than the M treatment, while a negative value denotes M > E. A strongly supported treatment effect is implied by most of the posterior lying to one side of zero. The vertical bars show the 66% and 95% quantiles and the median of the posterior distribution. This plot was created using posterior predictions of the means that were not adjusted for differences in dry weight between treatments.

Finally, we investigated the treatment × sex interaction term by calculating the posterior difference in the treatment effect size between sexes. Though none of the results reached statistical significance, there was some indication that the decrease in carbohydrate content (in E relative to M) was greater in males than females (PP = 0.092), and that the sexes showed different changes in protein content (PP = 0.11) (Fig. S6).

## DISCUSSION

We found that experimental manipulation of sexual selection and sexual conflict caused several physiological and life history traits to evolve. First, flies from the elevated polyandry (E) treatment took longer to develop into adults than their monogamous (M) counterparts and the difference in development time between the sexes was greater in the E lines. Second, E treatment flies of both sexes had faster metabolic rates, largely due to E line flies being more active than M line flies. Third, macro-metabolite composition has diverged between sexual selection treatments, such that E females had a greater lipid content than M females, E males greater glycogen content than M males, and E males less chitin and sugar content than M males. Finally, E males were less resistant to desiccation and starvation than were M males, whereas E females were more resistant than M females.

E males produce a faster and more vigorous courtship song (Debelle *et al*., 2014, 2017), more cuticular hydrocarbons (Hunt *et al*., 2012), and larger accessory glands (Crudgington *et al*., 2009) than M males. These sexually selected traits may be energetically costly to produce and maintain, demanding greater metabolic activity (Montooth *et al*., 2003; Immonen *et al*., 2016; Berson *et al*., 2019). Metabolic rate showed a sex-specific response to selection for early life reproduction in bean beetles (Arnqvist *et al*., 2017). However, locomotor activity and metabolic rate have a significant intersexual genetic correlation in *D. melanogaster* (Long & Rice, 2007; Nandy *et al*., 2013; Videlier *et al*., 2021). Therefore, sexual selection favouring increased activity and metabolic rate in males may generate sexual conflict by increasing activity and metabolic rate above optimal levels for females (Hämäläinen *et al*., 2018). We found no difference between treatments or sexes in the respiratory quotient (RQ), which describes metabolic substrate use, despite evolved differences in activity and metabolic rate. The lack of evolution of RQ may suggest that E flies have evolved compensatory physiological mechanisms to offset the metabolic costs of increased energy expenditure (Husak & Swallow, 2011).

We found that E males had higher glycogen content than M males. Carbohydrates are the main fuel used during intense aerobic activities, such as flight (Wigglesworth, 1949) and courtship (Bertram *et al*., 2011), and glycogen provides the main source of trehalose marshalled during intense activity (Becker *et al*., 1996). Sexual selection favouring endurance capacity may also increase lipid respiration and fat storage (Gyulavári *et al*., 2014). However, despite male courtship song being an endurance contest (Debelle *et al*., 2017), in males we found no difference between treatments in lipid content. Limited time and resources may demand strategic resource allocation decisions. We did find higher lipid content in E females, possibly reflecting selection for endurance capacity, as E females are courted more frequently (Crudgington *et al*., 2005; Debelle *et al*., 2014). No metabolites showed a significant treatment × sex interaction, indicating the response to sexual selection treatment did not differ between the sexes, potentially highlighting a strong intersexual genetic correlation constraining divergence of physiological traits (Videlier *et al*., 2021; Wittman *et al*., 2021). Our results suggest that heightened sexual selection and sexual conflict favours storage and regulation of metabolites to meet increased metabolic demands (Montooth *et al*., 2003; Crudgington *et al*., 2009; Debelle *et al*., 2014, 2017; Gyulavári *et al*., 2014), whereas relaxed sexual selection may instead enable investment in other components of fitness.

E males were less tolerant of desiccation and starvation than M males despite their greater glycogen content, which buffers against these stresses (Marron *et al*., 2003). Previous work artificially selecting on desiccation tolerance in *D. melanogaster* found selected lines had lower metabolic rates (Hoffmann & Parsons, 1989); our experiment imposed different selective pressures but found the same negative correlation between metabolic rate and stress resistance. The E lines have greater cuticular hydrocarbon (CH) content but there is no difference in the abundance of long-chain CHs (which offer greater desiccation resistance) between treatments (Hunt *et al*., 2012). That E males are less stress resistant despite having more macromolecules which buffer against such stressors suggests a trade-off, where heightened sexual selection favours investment in current reproduction at a cost to later-life survival, while the relaxation of sexual selection favours increased investment in longevity (Kotiaho, 2001; Hunt *et al*., 2004; Nandy *et al*., 2013). In contrast, E females showed no such trade-off, as E females were as stress resistant as M females, despite investing more in fecundity (Crudgington *et al*., 2005; Immonen *et al*., 2014). The greater lipid content we found in E females could mitigate such a trade-off, because lipids provide protection against starvation alongside being a major component of eggs (Chippindale *et al*., 1996). Thus, the sexes have altered investment in macro-metabolite composition under sexual selection, which may allow differential investment in sexually selected traits and sexual conflict persistence and resistance traits. The subsequent fitness consequences during stress have different outcomes on the sexes. This suggests sex-specific selection in the adult stage which could further generate sexual conflict over shared traits, particularly during the juvenile stage (Badyaev, 2002).

Related to the juvenile stage, in both sexes, E flies took significantly longer to reach adulthood than flies evolving under monogamy, perhaps due to pleiotropic costs of increased investment in traits involved in sexual selection or sexual conflict (Zera & Harshman, 2001; Simmons *et al*., 2017; Berson *et al*., 2019). Moreover, because development time is probably genetically correlated across sexes (Lewis *et al*., 2011; Berger *et al*., 2014), the fact that both sexes evolved longer development might reflect intralocus sexual conflict. For example, E males might have evolved to invest more in sexual traits, slowing their development and causing a similar delay in E females because of pleiotropy. This would be similar to the results of Harano et al. (Harano *et al*., 2010), who selected for male beetles with larger mandibles (a sexually-selected trait), which caused maladaptive morphological evolution in females despite females lacking the male trait that was under selection (Harano *et al*., 2010). Conversely, longer development might benefit females, e.g. by enabling increased resource acquisition and allocation towards reproduction or resistance to male harm (Crudgington *et al*., 2005, 2010; Immonen *et al*., 2014). The sex difference in development time was also greater in the E lines, possibly resulting from evolution towards the (shorter) female optimum under monogamy.

To conclude, we have shown that experimental manipulation of sexual selection and sexual conflict caused the evolution of several traits that are not directly involved in sexual interactions. Differences in activity levels and metabolic rate coincided with differential investment in macro-metabolites, reflecting the energetic demands associated with the elevation or removal of sexual selection. Males evolving under elevated polyandry suffered reduced survival, potentially trading off current vs. future reproduction. However, the same was not true of females, perhaps due to selection favouring endurance capacity in females to offset the costs of frequent male courtship. The physiological and life history traits we measured likely have a shared genetic basis, and thus may be subject to intralocus sexual conflict. The magnitude of any sexual dimorphism did not differ between treatments for the physiological traits we measured (macro-metabolites and metabolic rate), whereas life history traits (development time and stress resistance) were more dimorphic in populations evolving under elevated polyandry, suggesting tighter intersexual genetic correlations over physiological traits. Overall, our findings highlight that sexual selection results in coordinated evolution of fundamental physiological and non-reproductive life history traits, implicating sexual selection as an important factor in the evolution of life history strategies.

## ACKNOWLEDGEMENTS

We would like to thank Göran Arnqvist who kindly offered the use of his laboratory to measure metabolic rates. Andrew Beckerman, Dylan Childs, John Jackson and Björn Rogell provided helpful discussion about statistical analysis and Henry Barton helped relabel images for blind analysis. Many members of the Snook lab contributed to running the experimental sexual selection lines over the years. This work was funded by the National Environment Research Council (NERC) through the Adapting to the Challenges of a Changing Environment Doctoral Training Partnership (ACCE DTP; NE/L002450/1). Previous funding for the lines came from: the US National Science Foundation (DEB-0093149) and NERC grants (NE/B504065/1, NE/D003741/1, NE/I014632/1).

## CONTRIBUTIONS

RRS designed the experiments. ALB, MDG, ZKN, and RRS collected the data. MDG and LH performed statistical analyses. MDG, LH and RRS wrote the manuscript. All authors agreed to the final version of the manuscript.

## DATA AVAILABILITY STATEMENT

Data and code are available from the Dryad digital repository: https://doi.org/10.5061/dryad.9cnp5hqhk, and GitHub: https://lukeholman.github.io/exp_evol_respiration/

