## Supplementary methods and figures for "Experimental sexual selection affects the evolution of physiological and life history traits"

**ORCiD:** MDG: 0000-0002-3426-4341, LH: 0000-000-2-7268-2173, ALB: 0000-0002-2556-5249, ZKN: 0000-0001-5157-3229, RRS: 0000-0003-1852-1448.

Data and code are available from the Dryad digital repository: https://doi.org/10.5061/dryad.9cnp5hqhk, and GitHub: https://lukeholman.github.io/exp_evol_respiration/

### SUPPLEMENTAL METHODS

#### *Metabolite extractions*

We measured metabolite composition at generations 199, 198, 197 and 195 for lines 1-4, respectively. Triads (n = 3) were weighed to the nearest 1µg (METTLER TOLEDO® UMX2 ultra-microbalance) and flash frozen in liquid nitrogen. Triads were then placed in a 0.35ml glass vial insert (SUPELCO Analytical®) of known weight, dried at 55ºC overnight, and re-weighed to obtain dry weight.

*Lipids*

To extract lipids, 200µl of hexane (Fisher scientific®) was added to each sample, which was then vacuum infiltrated and incubated at room temperature overnight. The supernatant was discarded, and samples dried overnight at 55ºC. The lipid content was determined by subtracting the dry weight after hexane extraction from the initial dry weight. Note that the hexane fraction is a measure of total lipid content, including both metabolic reserves and structural lipids of cell membranes. We assume that structural lipids do not vary much between treatments.

*Soluble carbohydrates*

After hexane extraction, samples were placed in 200µl of 80% ethanol (Fisher scientific®), vacuum infiltrated, and incubated at room temperature overnight. The supernatant was discarded, and samples dried overnight at 55ºC. The soluble carbohydrate content was calculated as the dry weight after hexane extraction minus the dry weight after ethanol extraction.

*Soluble protein*

After ethanol extraction, dried samples were transferred to a screwcap tube (SUPELCO Analytical®) and ground before adding 200µl of Tris buffer (20mM, pH 7.0; Fisons Analytical Reagents®) and centrifuged at 16000G. A 10µl aliquot of supernatant from each sample was loaded on a 96-well plate containing 200µl bicinchoninic acid protein assay reagent (Bio-Rad®). Protein concentrations were determined using standards of bovine serum albumin (SIGMA-Aldrich®) at an absorbance of 562nm (FLUOstar OPTIMA® plate reader, BMG labtech).

*Glycogen*

Glycogen extraction protocol was modified from Caporn et al. (Caporn *et al.*, 1999). Remaining samples in Tris buffer were autoclaved before adding 100µl of MES (500mM, pH 4.5; SIGMA-Aldrich®), containing 4 units of a-amylase from *Aspergillus oryzae* (SIGMA-Aldrich®) and 14 units of Amylglucosidase from *A. niger* (SIGMA-Aldrich®) and incubated at 37ºC for 4 hours. Samples were centrifuged at 16000G and 50µl of supernatant loaded on a 96 well-plate containing 200µl of 100mM HEPES (pH 7.4; Roche®), 5mM magnesium chloride (Fisons Analytical Reagents®), 1.6mM NAD (SIGMA-Aldrich®), 4mM ATP, 0.5 U glucose-6-phospate dehydrogenase (Roche®). Glucose concentration was determined by the addition of 0.5 units of hexokinase (Roche®) taking readings at 340nm.

*Chitin*

After centrifugation in Tris buffer, the pellet was incubated at 100ºC for two hours with KOH (BDH Laboratory Reagents®) to remove enzyme contaminants, centrifuged again, and the pellet then washed three times in ddH20. The remaining residue was transferred to the original vial insert (used to obtain dry weight), dried and reweighed. Chitin content was determined by subtracting the final dry weight from the initial dry weight.

***Statistical analysis***

All statistical analyses were performed in R version 4.0.3 (R Core Team, 2020). We fit models using Bayesian approaches in *brms* (Bürkner, 2017), and specified regularising (i.e. conservative) priors on the fixed and random effects. Complete code and description of models are provided in the code repository [https://anonymous.4open.science/r/exp_evol_respiration-2647]. All models included replicate line as a random intercept to reflect the experimental design of the selection experiment (i.e. where n = 8). We also performed analyses with selection treatment fitted as a random slope to allow lines to vary in their response to selection (Schielzeth & Forstmeier, 2009), however results were qualitatively identical, and thus we present only the results of models without random slopes in the Results.

***Statistical analysis of development time and survival (survival analysis):*** We analysed juvenile development time using Cox proportional hazards survival analysis, where time (days elapsed since seeding day) to event (eclosion) was used as the response. Flies that did not eclose within the observation period were right censored on the last collection day. Sex was assigned to censored individuals by calculating the observed sex ratio of eclosees from each vial and assigning the appropriate sex ratio to the remaining uneclosed individuals of the 100 larvae initially seeded (assuming an equal 50:50 sex ratio of larvae). We included sexual selection treatment, sex, and the treatment x sex interaction as fixed effects and seeding day as a covariate. We included experimental evolution line as a random intercept term, with selection treatment as a random slope term. We also included vial ID as a random intercept term as larvae seeded in the same vial may show a correlated response.

We analysed wing length differences with sexual selection treatment, sex, and the treatment x sex interaction as fixed effects and seeding day as a covariate, and experimental evolution line as a random intercept term with a random slope for selection treatment.

For desiccation and starvation resistance preliminary analyses indicated violation of the proportional hazards assumption due to crossing hazards (see code repository). Therefore, we used accelerated failure time models with a Weibull distribution to model survival. Time (in hours) to event (death) was used as the response, with sexual selection treatment, sex, and the treatment x sex interaction as fixed effects. We included experimental evolution line as a random intercept term and a random slope for selection treatment. We also included vial ID as a random intercept term as individuals housed in the same vial may show a correlated response. For the starvation resistance assay, five individuals (two M females and three E females) were right censored at the end of the observation period as death times were not recorded (including one fly which remained alive).

**Statistical analysis of respiration:** Graphical exploration of the respirometry data revealed that activity level (and to a lesser extent body weight) was correlated with respiration, and also varied by sex and treatment (Fig. S2). We therefore analysed these data using a structural equation model (SEM), motivated by the causal relationships that we hypothesise to exist between the variables, given the experimental design and our biological understanding of the system (Figure S1A). The model treats activity level and body weight as ‘mediator variables’, which potentially mediate some of the effect of predictors such as treatment and sex on respiration. Furthermore, $VO_2$ and $VCO_2$ were highly correlated, and so we focused the analysis on just one of these (namely $VO_2$), and also searched for predictors of the respiratory quotient RQ (i.e., $VCO_2$ / $VO_2$). We did not perform model selection to ‘prune out’ causal relationships for which there was no evidence, both for philosophical and computational reasons; rather, we carefully constructed a model that we believe to be biologically plausible and then estimated its parameters (some of which differ from zero, providing evidence for correlations or causal relationships). The SEM contained the following submodels (in *brms*-like pseudocode):

$VCO_2$ ~ $O_2$ × RQ
RQ ~ (0.7 + 0.3 × *inverse_logit*($RQ_{est}$))
$VO_2$ ~ Treatment x Sex x Cycle + Activity + Weight + (Treatment |p| Line) + (1 | Triad)
$RQ_{est}$ ~ Treatment x Sex x Cycle + Activity + Weight + (Treatment |p| Line) + (1 | Triad)
Activity ~ Treatment x Sex x Cycle + (Treatment |p| Line) + (1 | Triad)
Weight ~ Treatment x Sex + (Treatment |p| Line)

Here, multiplication signs denote two- and three-way interactions among the fixed effects, random effects are feature parentheses and the pipe symbol, and *inverse_logit* denotes the inverse logit function being applied to an internally-estimated parameter $RQ_{est}$ (which is estimated on the link scale, i.e. -∞ to ∞). Note that we estimated effects on $VO_2$ and $RQ_{est}$, but not directly to $VCO_2$; $VCO_2$ is instead predicted as $VO_2$ x RQ, where RQ is calculated from $RQ_{est}$ using a modified inverse logit function which backtransforms from the link scale to a value in the range 0.7 - 1, which is the true possible range of RQ (which is determined by the chemical equations of respiration and the ratios of metabolic substrates). Furthermore, activity level and body weight have their own sub-models (allowing them to vary by sex, treatment, line, and triad), but they are also predictor variables of $VO_2$ and RQ, allowing them to ‘mediate’ some of the effects of sex, treatment, line, and triad on $VO_2$ and RQ. Regarding the random effects, the *brms* notation indicates that the model fits a random intercept for Line and Triad, as well as a random slope for treatment (allowing the replicate lines to differ in their response to treatment). The notation |p| indicates that the SEM estimates and accounts for correlations in the line effects between the response variables; for example, the model tests whether lines with larger body mass are also more active, and accounts for this correlation when estimating the fixed effects. After fitting the model, we calculated the posterior estimates of the means for the mediator variables, and the treatment effect sizes (Cohen’s *d*) for $VO_2$ and RQ, both with and without the moderating effects of activity and weight.

**Statistical analysis of metabolite composition:**

We began by expressing the abundance of each metabolite as a proportion of dry weight and scaling each response variable to have mean zero and unit standard deviation. Graphical exploration of the metabolite data revealed that some of the metabolite measurements were inter-correlated, some were correlated with dry body weight, and dry mass differed between sexes and the M and E treatments (Fig. S4). We therefore analysed metabolites using a structural equation model (SEM), motivated by the hypothesised causal diagram shown in Fig. S1B, which treats dry weight as a 'mediator variable' that mediates some of the effect of treatment and sex on each metabolite.

In this SEM, the sub-model for dry weight included treatment, sex, and their interaction as fixed effects, and included 'line' as a random intercept; we also allowed for differences in the treatment effect between lines (modelled as random slopes). The sub-models for each of the five metabolites were similar, but they included dry weight as an additional covariate. After fitting the model, we calculated the treatment effect size (Cohen's *d*) for each metabolite, both with and without the moderating effect of dry weight. We also calculated the difference in treatment effect size between the sexes, to test for a treatment x sex interaction.

### RESULTS

Table S1. Respirometry SEM table. This tables shows the fixed effects estimates for treatment, sex, their interaction, as well as the slope associated with dry weight (where relevant), for each of the response variables. The `p` column shows 1 - minus the "probability of direction", i.e., the posterior probability that the reported sign of the estimate is correct given the data and the prior; subtracting this value from one gives a Bayesian equivalent of a one-sided p-value. For brevity, we have omitted the estimates of residual (co)variance.

| | Parameter | Estimate | Est.Error | Q2.5 | Q97.5 | p |
|---|---|---|---|---|---|---|
| Oxygen consumption (VO2) | **Intercept** | **8.057** | **0.414** | **7.242** | **8.865** | **< 0.001** |
| | Treatment (E) | -0.135 | 0.516 | -1.146 | 0.890 | 0.395 |
| | **Sex (M)** | **-1.030** | **0.499** | **-2.005** | **-0.039** | **0.021** |
| | **Cycle (II)** | **-1.102** | **0.336** | **-1.755** | **-0.439** | **< 0.001** |
| | **Cycle (III)** | **-1.726** | **0.329** | **-2.359** | **-1.069** | **< 0.001** |
| | Body weight | 0.275 | 0.233 | -0.179 | 0.737 | 0.119 |
| | **Activity level** | **1.015** | **0.167** | **0.688** | **1.342** | **< 0.001** |
| | Treatment (E) x Sex (M) | 0.635 | 0.575 | -0.508 | 1.749 | 0.136 |
| | Treatment (E) x Cycle (II) | 0.305 | 0.447 | -0.570 | 1.163 | 0.248 |
| | Treatment (E) x Cycle (III) | -0.138 | 0.433 | -0.990 | 0.710 | 0.377 |
| | Sex (M) x Cycle (II) | 0.152 | 0.439 | -0.707 | 1.000 | 0.358 |
| | Sex (M) x Cycle (III) | -0.507 | 0.445 | -1.382 | 0.368 | 0.123 |
| | Treatment (E) x Sex (M) x Cycle (II) | -0.161 | 0.573 | -1.273 | 0.961 | 0.388 |
| | Treatment (E) x Sex (M) x Cycle (III) | 0.509 | 0.562 | -0.596 | 1.618 | 0.182 |
| Respiratory quotient (RQ) | Intercept | 0.000 | 0.316 | -0.600 | 0.655 | 0.493 |
| | Treatment (E) | 0.286 | 0.436 | -0.596 | 1.137 | 0.247 |

| | | | | | | |
|---|---|---|---|---|---|---|
| | Sex (M) | 0.467 | 0.423 | -0.349 | 1.303 | 0.131 |
| | Cycle (II) | 0.378 | 0.375 | -0.360 | 1.123 | 0.155 |
| | Cycle (III) | -0.107 | 0.378 | -0.839 | 0.653 | 0.385 |
| | Body weight | -0.014 | 0.181 | -0.372 | 0.348 | 0.463 |
| | Activity level | -0.218 | 0.155 | -0.517 | 0.095 | 0.079 |
| | Treatment (E) x Sex (M) | 0.140 | 0.501 | -0.868 | 1.105 | 0.385 |
| | Treatment (E) x Cycle (II) | 0.050 | 0.463 | -0.847 | 0.972 | 0.456 |
| | Treatment (E) x Cycle (III) | 0.565 | 0.481 | -0.391 | 1.524 | 0.116 |
| | Sex (M) x Cycle (II) | -0.507 | 0.528 | -1.544 | 0.542 | 0.166 |
| | Sex (M) x Cycle (III) | -0.031 | 0.579 | -1.142 | 1.120 | 0.474 |
| | Treatment (E) x Sex (M) x Cycle (II) | -0.483 | 0.608 | -1.684 | 0.698 | 0.212 |
| | Treatment (E) x Sex (M) x Cycle (III) | -0.805 | 0.664 | -2.109 | 0.494 | 0.111 |
| Activity level | **Intercept** | **-0.784** | **0.223** | **-1.225** | **-0.343** | **0.001** |
| | **Treatment (E)** | **1.060** | **0.303** | **0.467** | **1.650** | **0.001** |
| | Sex (M) | 0.467 | 0.283 | -0.096 | 1.012 | 0.050 |
| | Cycle (II) | 0.196 | 0.184 | -0.169 | 0.557 | 0.141 |
| | Cycle (III) | 0.243 | 0.187 | -0.122 | 0.608 | 0.097 |
| | Treatment (E) x Sex (M) | -0.072 | 0.380 | -0.826 | 0.675 | 0.424 |
| | Treatment (E) x Cycle (II) | 0.099 | 0.256 | -0.399 | 0.607 | 0.346 |
| | Treatment (E) x Cycle (III) | -0.016 | 0.256 | -0.517 | 0.482 | 0.474 |
| | **Sex (M) x Cycle (II)** | **-0.485** | **0.250** | **-0.978** | **0.005** | **0.026** |
| | **Sex (M) x Cycle (III)** | **-0.683** | **0.258** | **-1.187** | **-0.174** | **0.005** |
| | Treatment (E) x Sex (M) x Cycle (II) | 0.333 | 0.351 | -0.348 | 1.022 | 0.171 |
| | Treatment (E) x Sex (M) x Cycle (III) | 0.529 | 0.352 | -0.162 | 1.215 | 0.067 |
| Body weight | Intercept | 0.361 | 0.276 | -0.179 | 0.913 | 0.086 |
| | Treatment (E) | 0.518 | 0.367 | -0.229 | 1.218 | 0.075 |
| | **Sex (M)** | **-1.022** | **0.258** | **-1.525** | **-0.516** | **< 0.001** |
| | Treatment (E) x Sex (M) | -0.439 | 0.353 | -1.132 | 0.262 | 0.105 |

Table S2. Metabolite SEM table. This tables shows the fixed effects estimates for treatment, sex, their interaction, as well as the slope associated with dry weight (where relevant), for each of the six response variables. The `p` column shows 1 - minus the "probability of direction", i.e., the posterior probability that the reported sign of the estimate is correct given the data and the prior; subtracting this value from one gives a Bayesian equivalent of a one-sided p-value. For brevity, we have omitted the estimates of residual (co)variance.

| | Parameter | Estimate | Est. error | CI lower | CI upper | p |
|---|---|---|---|---|---|---|
| Carbohydrates | Dry weight | 0.105 | 0.269 | -0.419 | 0.622 | 0.352 |
| | Sex (M) | 0.024 | 0.427 | -0.812 | 0.864 | 0.477 |
| | Treatment (E) | -0.246 | 0.302 | -0.832 | 0.348 | 0.207 |
| | Sex (M) x Treatment (E) | -0.414 | 0.347 | -1.09 | 0.263 | 0.117 |
| Chitin | **Dry weight** | **-0.486** | **0.26** | **-0.995** | **0.023** | **0.031** |
| | Sex (M) | 0.399 | 0.42 | -0.435 | 1.206 | 0.170 |
| | Treatment (E) | -0.113 | 0.285 | -0.673 | 0.453 | 0.341 |
| | Sex (M) x Treatment (E) | -0.317 | 0.328 | -0.957 | 0.32 | 0.166 |
| Glycogen | Dry weight | 0.332 | 0.267 | -0.18 | 0.85 | 0.110 |
| | Sex (M) | -0.267 | 0.427 | -1.096 | 0.567 | 0.265 |
| | Treatment (E) | 0.22 | 0.296 | -0.367 | 0.787 | 0.224 |
| | Sex (M) x Treatment (E) | 0.387 | 0.351 | -0.297 | 1.066 | 0.139 |
| Lipids | **Dry weight** | **0.539** | **0.254** | **0.041** | **1.029** | **0.017** |
| | Sex (M) | -0.119 | 0.411 | -0.926 | 0.68 | 0.382 |
| | Treatment (E) | 0.389 | 0.276 | -0.156 | 0.919 | 0.077 |
| | Sex (M) x Treatment (E) | -0.05 | 0.313 | -0.672 | 0.555 | 0.437 |
| Protein | Dry weight | -0.206 | 0.275 | -0.742 | 0.34 | 0.225 |
| | Sex (M) | -0.011 | 0.435 | -0.858 | 0.85 | 0.484 |
| | Treatment (E) | -0.216 | 0.306 | -0.818 | 0.378 | 0.241 |
| | Sex (M) x Treatment (E) | 0.352 | 0.361 | -0.347 | 1.057 | 0.167 |
| Dry weight | **Sex (M)** | **-1.614** | **0.143** | **-1.895** | **-1.334** | **< 0.001** |
| | **Treatment (E)** | **0.525** | **0.155** | **0.218** | **0.827** | **< 0.001** |
| | **Sex (M) x Treatment (E)** | **-0.354** | **0.196** | **-0.733** | **0.035** | **0.038** |

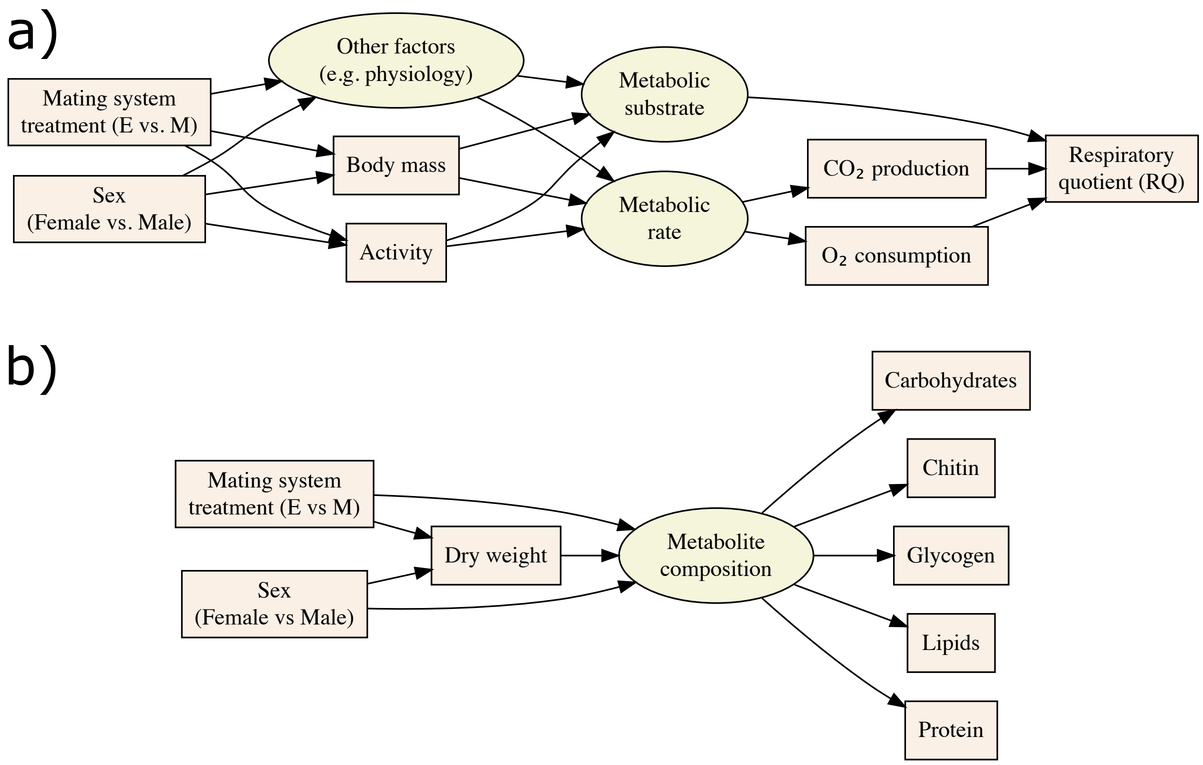


**<u>Figure S1.</u>** Directed acyclic graph (DAG) illustrates the causal pathways that we observed between the experimental or measured variables (square boxes) and latent variables (ovals). A) Metabolic rate. We hypothesise that sex and mating system potentially influence activity level and dry weight as well as other (e.g. physiological) factors. In turn, these variables may influence metabolic rate and metabolic substrate use. CO2 production and O2 consumption were measured as proxy for metabolic rate, and the respiratory quotient (RQ, ratio of CO2/O2) as a measure of metabolic substrate use. B) Macro-metabolite composition. We hypothesise that sex and mating system potentially influence dry weight as well as the metabolite composition (which we assessed by estimating the amount of carbohydrates, chitin, glycogen, lipids and protein). Additionally, dry weight is likely correlated with metabolite composition, and so dry weight acts as a 'mediator variable' between metabolite composition, and sex and treatment. The structural equation models described in the extended methods were built with these DAGs in mind.

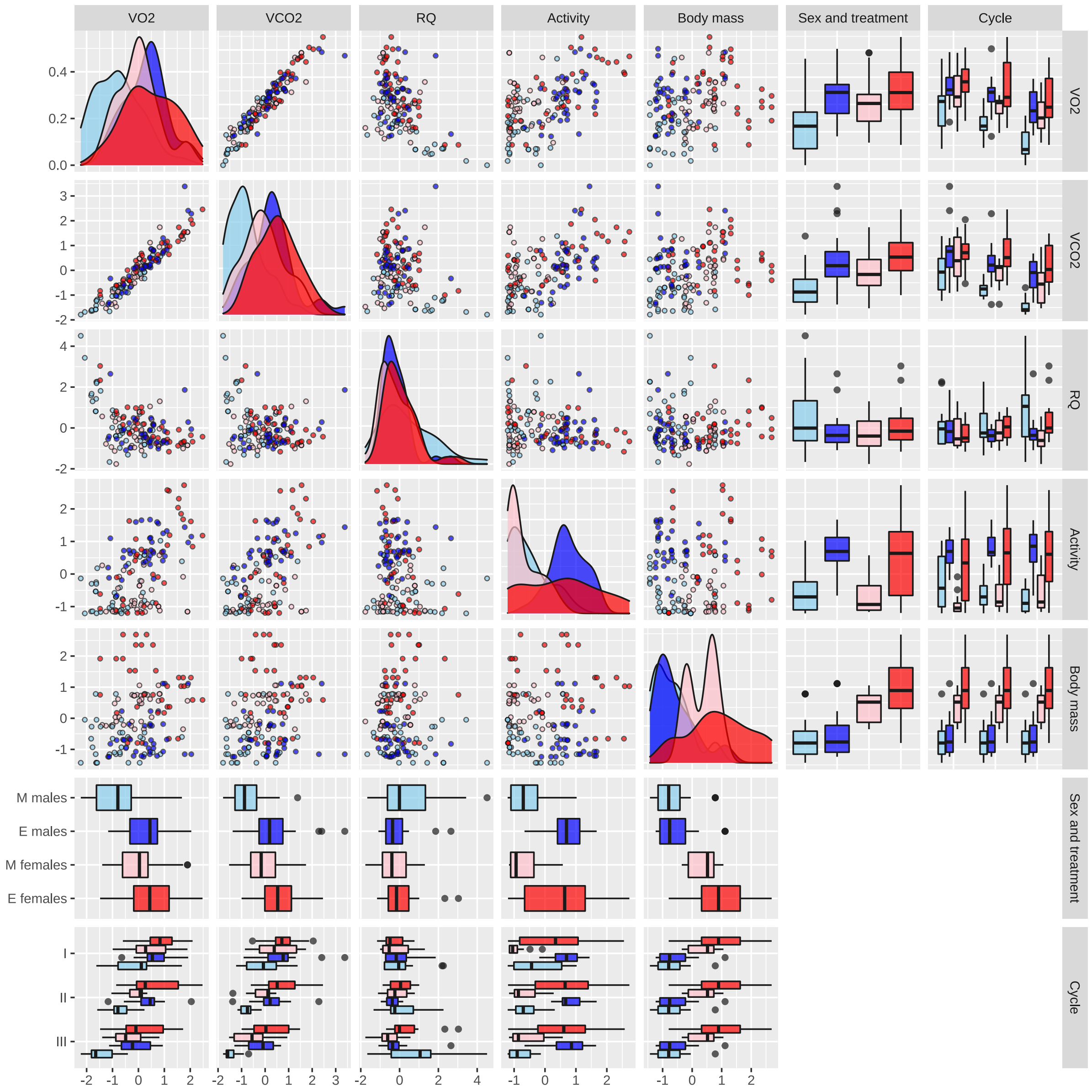


**<u>Figure S2</u>**: Respirometry raw data. Boxplots, scatterplots, and density plots illustrating the variance and covariance among the explanatory variables (Treatment, Sex, and Cycle), two mediator variables (Body mass and Activity), and three response variables ($O_2$, $CO_2$, and RQ). The data are split and coloured by treatment (red for females, blue for males). The plot shows the raw data in their original units, namely $mm^3$ of $O_2$ consumed or $CO_2$ produced the % time spent active, and the mass of the fly in milligrams (RQ is a ratio and thus has no units). Note that RQ is theoretically expected to lie in the range 0.7-1.0 because of the chemistry of respiration, but values outside this range often occur due to measurement error for $O_2$ and/or $CO_2$. Each point represents the measurement from a triad of same-sex, same-selection line flies.

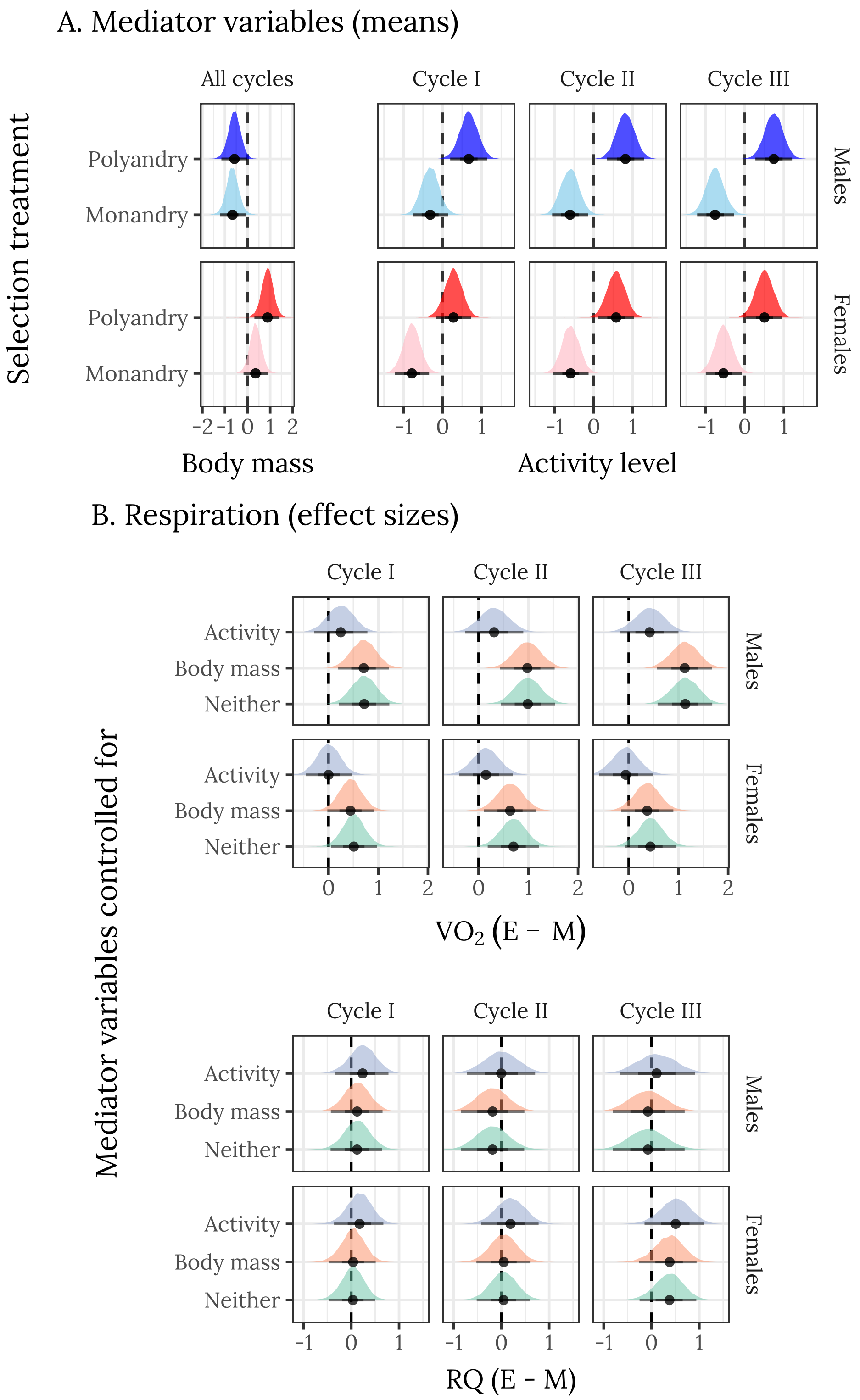


**<u>Figure S3.</u>** Effects of sex and selection treatment on metabolic rates and mediator variables (all cycles). A) Posterior estimates of the means for mediator variables (body mass and activity). Note that females are larger than males, and that E females are somewhat larger than M females. E individuals of both sexes are more active than M individuals, and there are changes in activity level across cycles (in particular, M males become less active with time, while P males maintain consistent, high activity levels). B) Posterior estimates of effect sizes of selection treatment (E - M) controlling for activity, body mass, or neither mediator variable. This was accomplished by producing the posterior predictions with activity or body mass set to its global mean value (i.e., 0), which is equivalent to asking what the response variables would be if the mediator variable were the same across sexes and treatments. Thus, the x-axes show the posterior estimate of standardised effect size (Cohen's d), i.e., a value of 1 would mean that the E treatment has a mean that is larger by 1 standard deviation. The thicker inner bars show the 66% quantiles of the posterior distribution, the thin outer bars show the 95% quantiles, and the points show the median. All response variables have been mean-centred and divided by the overall standard deviation to facilitate plotting (such that the dashed line at zero marks the mean across sexes, treatments and cycles).

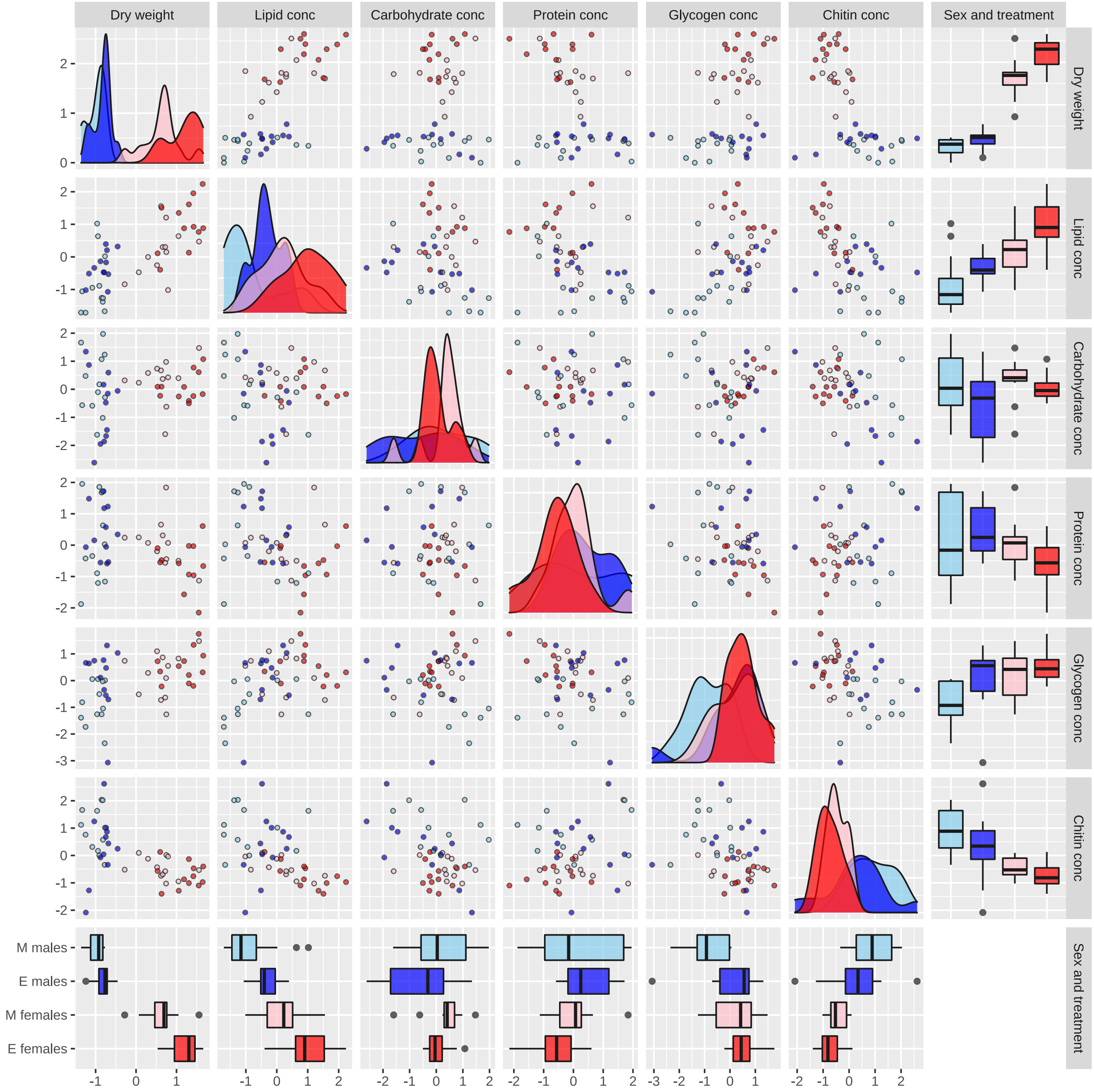


**Figure S4.** Metabolite raw data. Boxplots, scatterplots, and density plots illustrating the variance and covariance among the explanatory variables (Treatment, and Sex), mediator variable (Body mass), and five response variables (Lipid conc., Carbohydrate conc., Protein conc., Glycogen conc. Chitin conc.). Each measurement was calculated by dividing each metabolite by the dry weight of the triad of flies. Each point represents the measurement from a triad of same-sex, same-selection line flies. The data are split and coloured by treatment (red for females, blue for males).

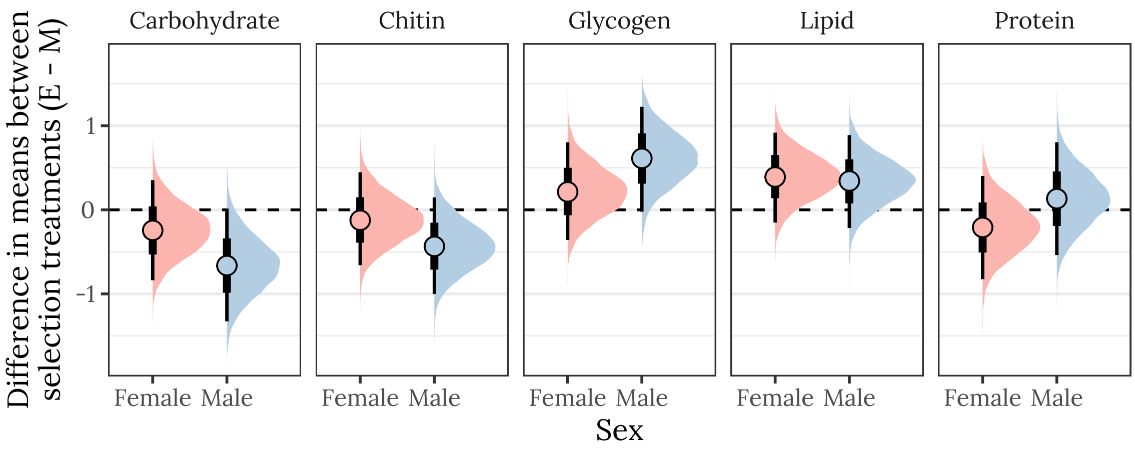


**<u>Figure S5.</u>** Posterior estimates of the treatment effect size for both sexes, for each of the five metabolites. A positive value means that the mean metabolite content is higher in the E treatment than the M treatment, while a negative value denotes M > E. A strongly supported treatment effect is implied by the majority of the posterior lying to one side of zero. The thicker inner bars show the 66% quantiles of the posterior distribution, the thin outer bars show the 95% quantiles, and the points show the median. This plot was created using posterior predictions of the means that were adjusted for differences in dry weight between treatments.

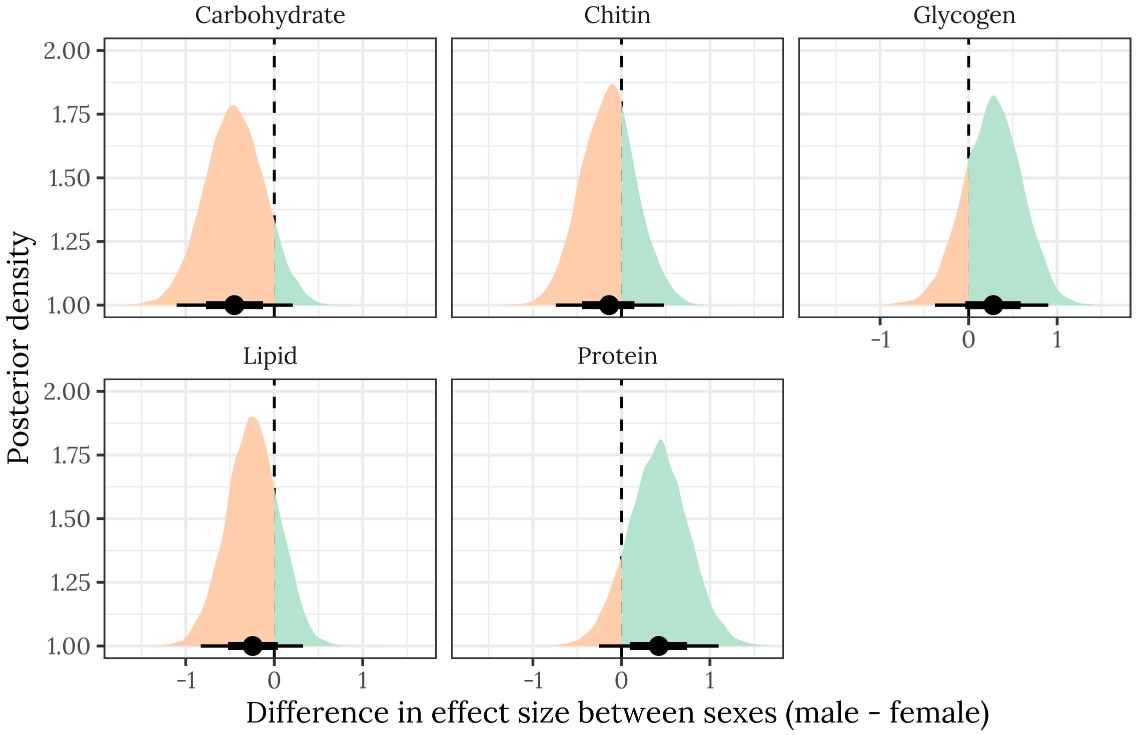


**<u>Figure S6.</u>** Posterior estimates of the difference in the treatment effect size (i.e., mean of E minus mean of M) between males and females, for each of the five metabolites. A positive value means that the effect size is more positive in males, and negative means it is more positive in females. A strongly supported sex difference in effect size would be implied by the majority of the posterior lying to one side of zero. The error bars summarise the 66% and 95% quantiles of the posterior. This plot was created using posterior predictions of the means that were not adjusted for differences in dry weight between treatments.
